# Sex-Specific Differential DNA Methylation in Mild Cognitive Impairment and Alzheimer’s Disease

**DOI:** 10.1101/2025.05.15.654279

**Authors:** Phillip E. Bergmann, Andy Madrid, Ligia A. Papale, Lindsay R. Clark, Sanjay Asthana, Sterling C. Johnson, Sündüz Keleş, Kirk J. Hogan, Reid S. Alisch

## Abstract

Sex differences in late-onset Alzheimer’s disease (AD) progression include accelerated decrements in cognitive status and greater amyloid and tau biomarker burdens in females. To identify sex-specific differentially methylated positions (DMPs) and genes in persons with mild cognitive impairment (MCI) and AD, we analyzed whole genome methylation sequencing on blood samples from participants with MCI (N=99, 52% female), AD (N=109, 43% female), and those cognitively unimpaired (CU; N=174, 52% female). Ninety-four percent of DMPs from MCI *vs*. CU, AD *vs*. CU, and AD *vs*. MCI pairwise comparisons were sex-specific. Female-specific DMPs were enriched in neurologic gene sets (*e.g*., synaptic membrane, ion channel complex), while male-specific DMPs showed limited enrichment. Sex-specific DMPs overlapped blood-specific enhancers, promoters, and transcription factor binding motifs, highlighting divergent epigenetic regulation by sex. These findings identify sex-specific genes and molecular pathways in MCI and AD and support that blood DNA methylation levels can distinguish cognitive status.

## Introduction

Late-onset dementia due to Alzheimer’s Disease (AD) occurs in 1 of 9 adults over the age of 65 years, with females accounting for nearly two thirds of diagnoses.^1^ Females experience accelerated trajectories of changes in cognitive status and elevated AD biomarker pathology compared to men.^2^ Mild cognitive impairment (MCI) is a prodromal stage of AD, with objective evidence of decrements in memory, language, and executive function with skills compatible with independent living. Eighty percent of persons with MCI progress to AD dementia within 6 years.^3^ Sex differences observed with conversion from CU to MCI are reported but are more subtle than with conversion from MCI to AD dementia.^4^ A steeper downward trajectory in cognitive performance and loss of hippocampal volume are superior predictors of MCI and AD in females than in males.^5,6^ Females convert from MCI to AD more rapidly with lower Mini-Mental State Examination scores than men at diagnosis.^7^ Aβ and tau biomarkers of neurodegeneration are more prevalent in females.^8^ In addition, the *APOE* ε4 genotype confers a heightened risk of developing AD in females than in males.^9^

The influences of environmental factors on AD pathogenesis are mediated in part by DNA methylation. Differential DNA methylation in blood correlates with the presence or absence of diverse neurodegenerative disorders^10^ and is seen in postmortem brain tissues from patients with AD.^11,12^ Patterns of differential DNA methylation that are shared between brain and blood^13^ support testing DNA methylation levels in accessible tissues with the potential for improved AD diagnosis and prognosis. We previously reported 9,756 differentially methylated positions (DMPs) between CU and MCI participants, and 14,530 DMPs between CU and AD participants.^14^ In the present investigation, we expand on these findings to test for DNA methylation levels that differ by sex between persons who are CU and those with MCI or AD.

## Results

### Participants

Whole genome methylation sequences in blood from 174 CU participants (52% female), 99 participants with MCI (52% female), and 109 participants with AD (43% female) were tested for differential methylation by cognitive status and sex (Table 1). Male participants were 2 to 4 years younger than female participants in each cognitive status group with 1 to 1.5 fewer years of education.

**Table 1:** Participant means, standard deviations, and demographic variables.

| Phenotype | Female |  |  | Male |  |  |
| --- | --- | --- | --- | --- | --- | --- |
|  | CU (N = 90) | AD (N = 47) | MCI (N = 51) | CU (N = 84) | AD (N = 62) | MCI (N = 48) |
| No *APOE* ε4 | 72 | 13 | 17 | 57 | 27 | 25 |
| Het. *APOE* ε4 | 15 | 23 | 25 | 25 | 25 | 19 |
| Hom. *APOE* ε4 | 3 | 11 | 9 | 1 | 9 | 3 |
| Unknown *APOE* ε4 status | 0 | 0 | 0 | 1 | 1 | 1 |
| Age (years) | 72.62 (5.03) | 74.04 (9.37) | 72.66 (7.19) | 66.87 (8.55) | 72.8 (10.06) | 75.56 (7.91) |
| Education (years) | 16.03 (2.34) | 14.34 (2.52) | 14.78 (2.12) | 17.4 (2.65) | 15.42 (2.96) | 15.7 (3.03) |
| Average bmi | 28.76 (5.97) | 25.15 (5.69) | 28.54 (7.09) | 28.65 (4.15) | 27.91 (3.68) | 29.33 (6.5) |
| ADRC | 54 | 44 | 29 | 73 | 60 | 40 |
| WRAP | 36 | 3 | 22 | 11 | 2 | 8 |
| American Indian or Alaska Native | 0 | 0 | 1 | 1 | 0 | 0 |
| Black or African American | 4 | 2 | 4 | 3 | 4 | 6 |
| White | 86 | 45 | 46 | 80 | 58 | 42 |

### Whole genome DNA methylation detection

With an average of 561 million paired-end reads for each participant, 539 million sequence reads uniquely aligned to the human genome reference sequence (GRCh38.14 [hg38]) with an average genomic coverage of 53X and a mean of 38X after merging across strands for interrogation of 25,406,945 CpGs. Batch effects in WGMS data tested with the first 2 principal components accounted for 5.1% and 2.5% of total variability. No batch effects for participant cohort (WADRC or WRAP), cognitive status, sex, or self-reported ancestry were observed.^14,15^

### Sex-Specific Differentially Methylated Positions (DMPs) in Mild Cognitive Impairment and Alzheimer’s Disease

#### Sex-specific DMPs between MCI and CU

4,326 female-specific and 2,345 male-specific DMPs were identified between MCI and CU participants, with 130 DMPs shared between sexes (Figs. 1A and 1B, Supplemental Table 1). 12 of the 130 DMPs shared between sexes have opposing DNA methylation levels (*e.g.,* lower DNA methylation in females and higher DNA methylation in males), indicating that they are also sex-specific. The majority of MCI-associated DMPs were hypermethylated, 55% in females and 54.4% in male (*i.e.*, greater methylation in MCI compared to CU participants). MCI-associated DMPs were fewer in each genic region compared to the distribution of all CpG sites analyzed with the exception of intergenic regions and CpG islands (Supplemental Figs.1A and 1B). Sex differences within genic and CpG island regions were observed in 1-5kb upstream genic structures, CpG island shores, and the open sea.

**Figure 1:**
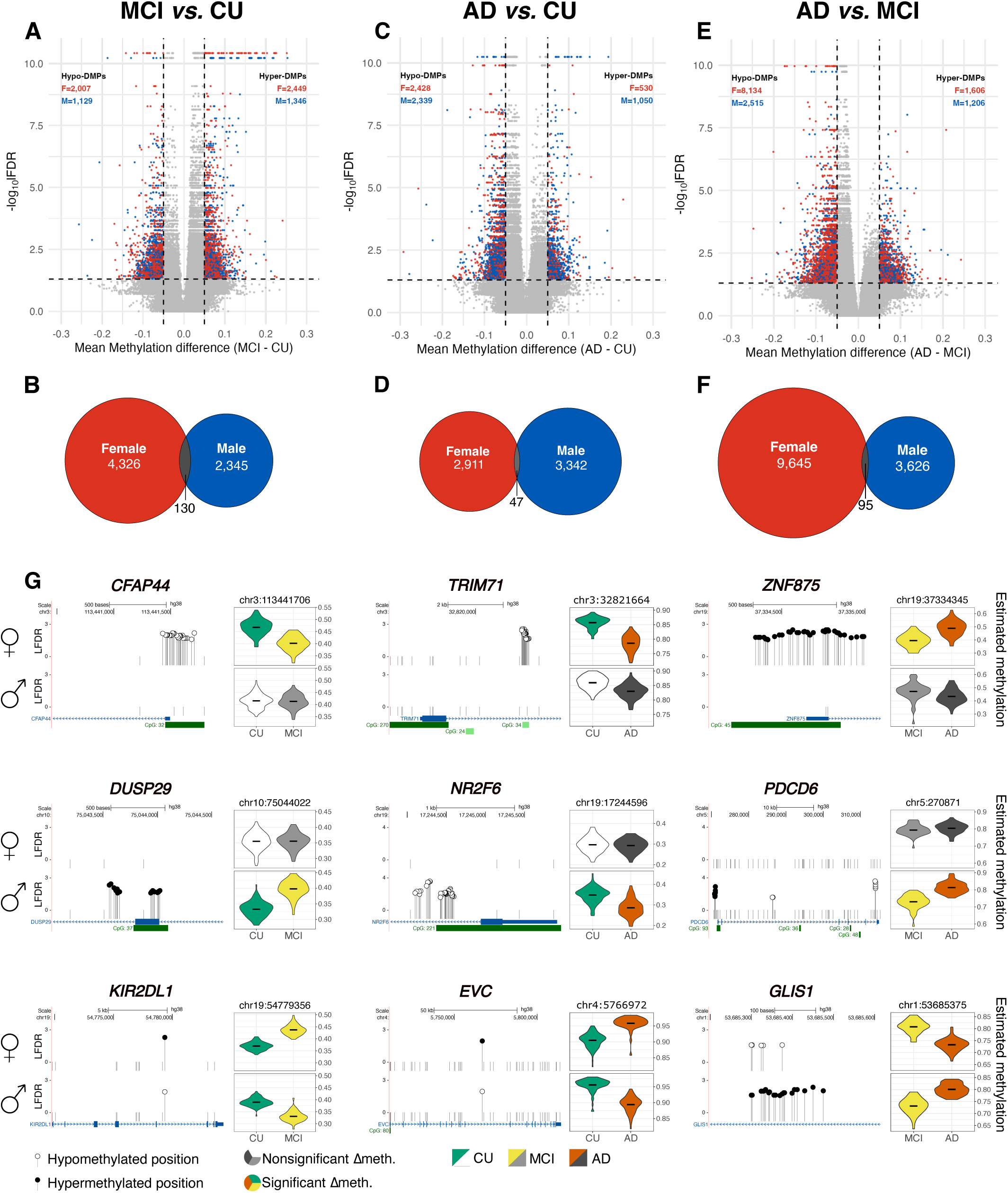
Identification of 5′-cytosine-phosphate-guanine-3′ (CpG) loci that comprise differentially methylated positions (DMPs) in blood between participants who are cognitively unimpaired (CU), those with mild cognitive impairment (MCI), and those with Alzheimer’s disease. Volcano plots of 25,244,219 WGMS CpG loci depicting the difference in mean methylation percent (x-axis, threshold > 5%) and the significance (y-axis, local false discovery rate [LFDR], threshold < 0.05) for (A) MCI *vs.* CU, (C) AD *vs.* CU, and (E) AD *vs.* MCI. Each point corresponds to a single CpG locus. Red points represent significant DMPs in females, blue points represent significant DMPs in males, and gray points indicate nonsignificant CpG loci (downsampled, 1:50). Venn diagrams depicting the specific and overlapping differentially methylated positions between females and males in the 3 pairwise comparisons (B) MCI *vs.* CU, (D) AD *vs.* CU, and (F) AD *vs.* MCI. (G) UCSC genome browser gene schematics depicting female-specific and male-specific DMPs, as well as shared DMPs between sexes with opposite methylation polarity between the pairwise comparisons. Sense and antisense strands are depicted with gene name abbreviations above each panel. Genomic distance (kb or bp) x-axis scales of each gene region are indicated at the top of each panel. Significance is indicated by local false discovery rate (LFDR, y-axis [-log_10_(LFDR)]), with longer stems and large white circles (hypomethylation) or large black circles (hypermethylation). Short stems extending only to zero without circles illustrate nonsignificant CpGs (1 out of 25 shown). Each gene diagram provides a track line for female (♀) DMPs (top panel) followed by (♂) male DMPs (bottom panel). The gene annotation track (blue, MANE Select Plus Clinical representative transcript) and the CpG island annotation track (green) are shown at the bottom of each panel. To the right of each gene diagram are two violin plots depicting the estimated methylation values in females (top) and males (bottom) of the indicated CpG locus (hg38 coordinates listed at top, 5’ most corresponding DMP in transcription orientation). Violin plots depicting the estimated methylation values (y-axis) in females and males of an example differentially methylated position (DMP, hg38 coordinates listed at top) that exhibits either significant (colored) or non-significant (grayscale) increments and decrements in DNA methylation levels by the diagnosis (*x*-axis). cognitively unimpaired (CU, green), mild cognitive impairment (MCI, yellow) and Alzheimer’s disease (AD, orange).

#### Sex-specific DMPs between AD and CU

2,911 female-specific and 3,342 male-specific DMPs were identified between AD and CU participants, with 47 DMPs shared between sexes (Figs. 1C and 1D, Supplemental Table 1). Nine of the 47 DMPs shared between sexes have opposing DNA methylation levels (*e.g.,* lower DNA methylation in females and higher DNA methylation in males), indicating that they are also sex specific. In contrast to the predominant hypermethylation observed in the sex-specific MCI-associated DMPs, the majority of sex-specific AD-associated DMPs were hypomethylated, 82.1% in females and 69% in males (*i.e.*, less methylation in AD compared to CU participants). AD-associated DMPs were fewer in each genic region compared to the distribution of all CpG sites analyzed, with the exception of intergenic regions and CpG islands (Supplemental Figs. 1C and 1D). Sex differences within genic and CpG island regions were observed in intergenic regions, 1-5kb upstream, 5’ UTRs, CpG islands, and the open sea.

#### Sex-specific DMPs between AD and MCI

9,645 female-specific and 3,628 male-specific DMPs were identified between AD and MCI participants, as well as 95 DMPs shared between sexes (Figs. 1E and 1F, Supplemental Table 1). 31 of the 95 DMPs shared between sexes have opposing DNA methylation levels (*e.g.,* lower DNA methylation in females and higher DNA methylation in males), indicating that they are also sex-specific. The majority of DMPs between MCI and AD were hypomethylated, 83.5% in females and 67.6% in males (*i.e.*, less methylation in participants with AD compared to MCI). These DMPs were fewer in each genic region and CpG islands compared to the distribution of all CpG sites analyzed (Supplemental Figs. 1E and 1F). Sex differences within genic regions and CpG islands were observed in 5’UTR regions, CpG islands, shores, and the open sea.

Clusters of more than one female-specific DMP for each pairwise comparison were observed in each of a subset of genes (Fig. 1G, Table 2) including *CFAP44* (MCI *vs.* CU), *TRIM71* (AD *vs.* CU), and *ZNF875* (AD *vs.* MCI). Clusters of more than one male-specific DMP for each pairwise comparison were observed in a subset of genes (Fig. 1G, Table 2) including *DUSP29* (MCI *vs.* CU), *NR2F6* (AD *vs.* CU), and *PDCD6* (AD *vs.* MCI). Certain DMPs were shared between the sexes in the 3 pairwise comparisons (Fig. 1G, Table 2) with opposite DNA methylation levels within a gene, including *KIR2DL1* (MCI *vs.* CU), *EVC* (AD *vs.* CU), and *GLIS1* (AD *vs.* MCI).

**Table 2:**
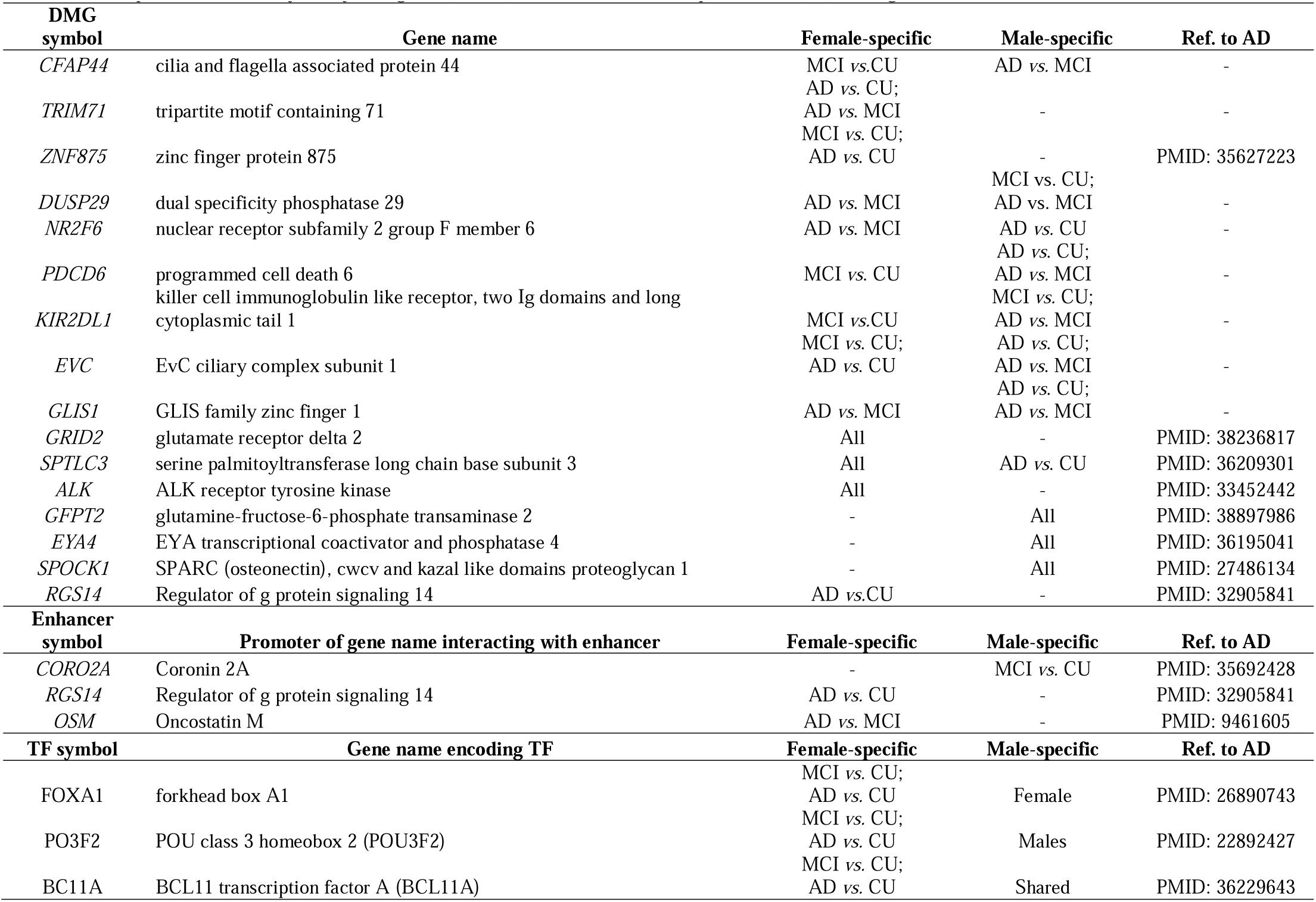
Sex-specific differentially methylated genes (DMGs), enhancer, and transcription factor (TF) binding motifs.

### Sex-specific Differentially Methylated Genes in Mild Cognitive Impairment and Alzheimer’s Disease

#### Sex-specific differentially methylated genes between MCI and CU

Annotation of the DMPs between participants with MCI and CU to genes identified 831 female-specific and 462 male-specific differentially methylated genes, as well as 157 genes differentially methylated in both sexes (Fig. 2A, Supplemental Table 2). Gene ontological enrichment analysis of the shared 157 differentially methylated genes yielded multiple neurological terms including regulation of trans-synaptic signaling, regulation of monoatomic ion transmembrane transport, and axonogenesis (Fig. 2B, Supplemental Table 3). Female-specific differentially methylated genes yielded neuronal terms including regulation of membrane potential, dendrite development, and synaptic transmission, glutamatergic (Fig. 2C). Male-specific differentially methylated genes between participants with MCI and CU were not enriched for any biological process term (Fig. 2D).

**Figure 2:**
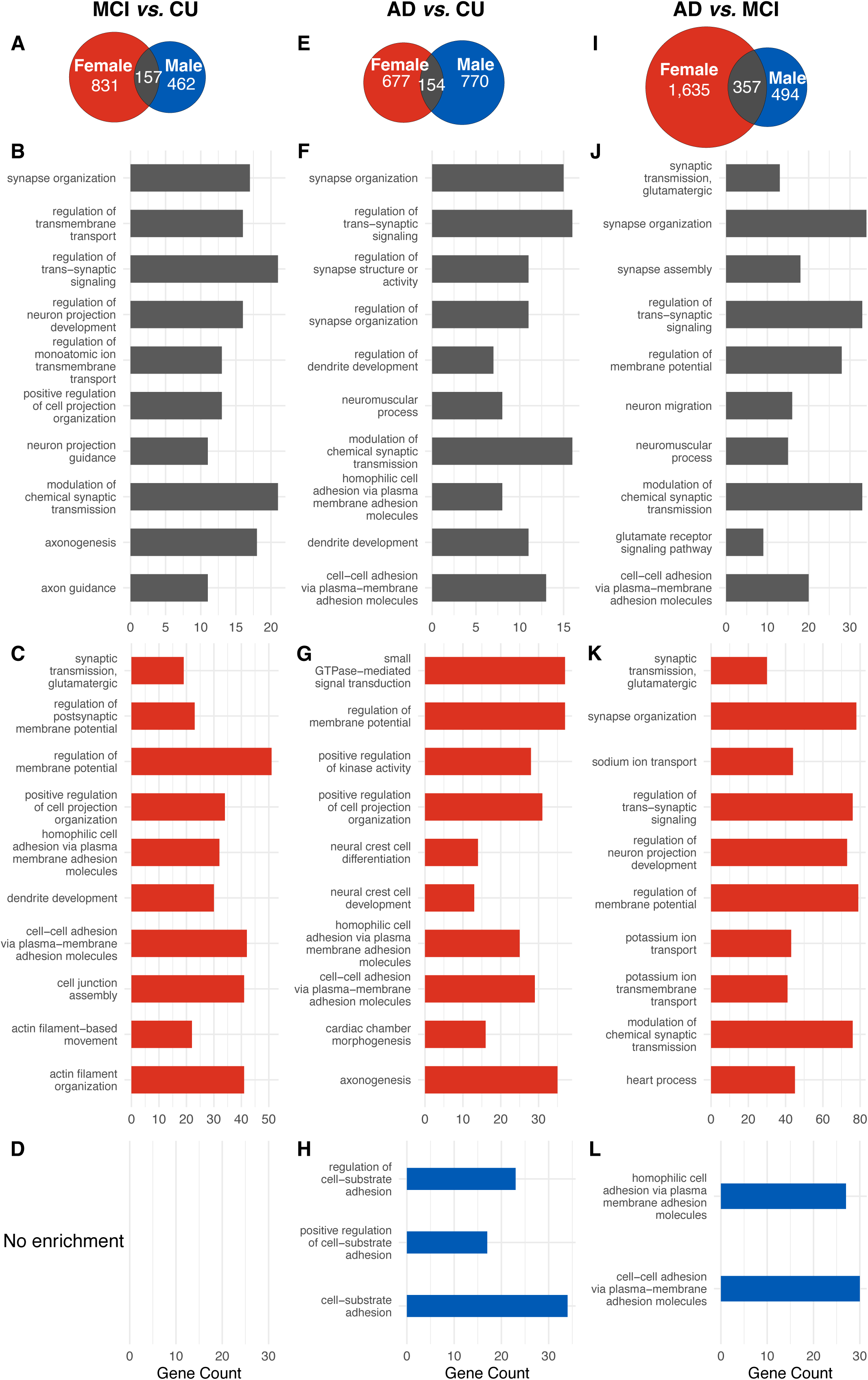
Sex-specific differentially methylated genes and enrichment of gene ontological biological processes across cognitive status comparisons. Venn diagrams depicting the sex-specific and overlapping differentially methylated genes between females and males in the 3 pairwise comparisons **(A)** MCI *vs.* CU, **(E)** AD *vs.* CU, and **(I)** AD *vs.* MCI. Barplots of enrichments of biological process gene ontological terms by adjusted *P*-value of shared [**(B)** MCI *vs.* CU, **(F)** AD *vs.* CU, and **(J)** AD *vs.* MCI], female-specific [**(C)** MCI *vs.* CU, **(G)** AD *vs.* CU, and **(K)** AD *vs.* MCI], and male-specific [**(D)** MCI *vs.* CU, **(H)** AD *vs.*CU, and **(L)** AD *vs.* MCI] differentially methylated genes for each pairwise comparison. The top 10 significant terms by significance are depicted (y-axis by false discovery rate < 0.05). The number of contributory genes (x-axis) associated with each term is provided together with the color-coordinated bars representing shared differentially methylated genes (gray), female-specific differentially methylated genes (red), and male-specific differentially methylated genes (blue).

#### Sex-specific differentially methylated genes between AD and CU

Annotation of the DMPs between participants with AD and CU to genes identified 677 female-specific and 770 male-specific differentially methylated genes, as well as 154 genes differentially methylated in both sexes (Fig. 2E, Supplemental Table 2). Gene ontological enrichment analysis of the 154 shared differentially methylated genes yielded multiple neurological terms and biological processes including modulation of chemical synaptic transmission, regulation of trans-synaptic signaling, and dendrite development (Fig. 2F, Supplemental Table 3). Female-specific differentially methylated genes yielded neuronal terms including regulation of membrane potential, neural crest cell development, and axonogenesis (Fig. 2G). Male-specific differentially methylated genes between participants with AD and CU were enriched for cellular cytoskeleton terms including cell-substrate adhesion (Fig. 2H).

#### Sex-specific differentially methylated genes between AD and MCI

Annotation of DMPs between participants with AD and MCI to genes identified 1,635 female-specific and 494 male-specific differentially methylated genes, as well as 357 genes differentially methylated in both sexes (Fig. 2I, Supplemental Table 2). Gene ontological enrichment analysis of the 357 differentially methylated genes shared between the sexes yielded multiple neurological terms including synapse organization, modulation of chemical synaptic transmission, and calcium ion transmembrane transport (Fig. 2J, Supplemental Table 3). Female-specific differentially methylated genes yielded neuronal terms including regulation of membrane potential, synapse organization, and regulation of postsynaptic membrane potential (Fig. 2K). Male-specific differentially methylated genes were enriched for plasma membrane adhesion terms (Fig. 2L).

### Sex-specific DMPs and differentially methylated genes shared between MCI *vs*. CU, AD *vs*. CU, and AD *vs*. MCI

13 DMPs in females and 14 DMPs in males were shared between the 3 sex-specific pairwise comparisons (Supplementary Figs. 2A and 2B). All 13 female-specific DMPs and 12 of 14 male-specific DMPs are located in an intergenic region near the centromere of chromosome 4, with 4 DMPs shared between males and females in all comparisons. Two male-specific DMPs are located at chromosome 1q12 more than 640 kilobases away from the nearest protein coding gene.

161 differentially methylated genes in females were shared between the 3 female pairwise comparisons, many with previously reported links to AD and neurodegeneration including *GRID2*, *SPTLC3*, and *ALK* (Figs. 3A, Table 2). 69 differentially methylated genes in males were shared in the 3 male pairwise comparisons, many with previously reported links to AD and neurodegeneration including *GFPT2*, *EYA4*, and *SPOCK1* (Fig. 3B, Table 2). 27 genes were differentially methylated in both females and males in the 6 pairwise comparisons (Fig. 3C). Thirty-seven female-specific differentially methylated genes shared between the 3 female pairwise comparisons were not differentially methylated in any male pairwise comparison. Six male-specific differentially methylated genes shared between the 3 male pairwise comparisons were not differentially methylated in any female comparisons.

**Figure 3:**
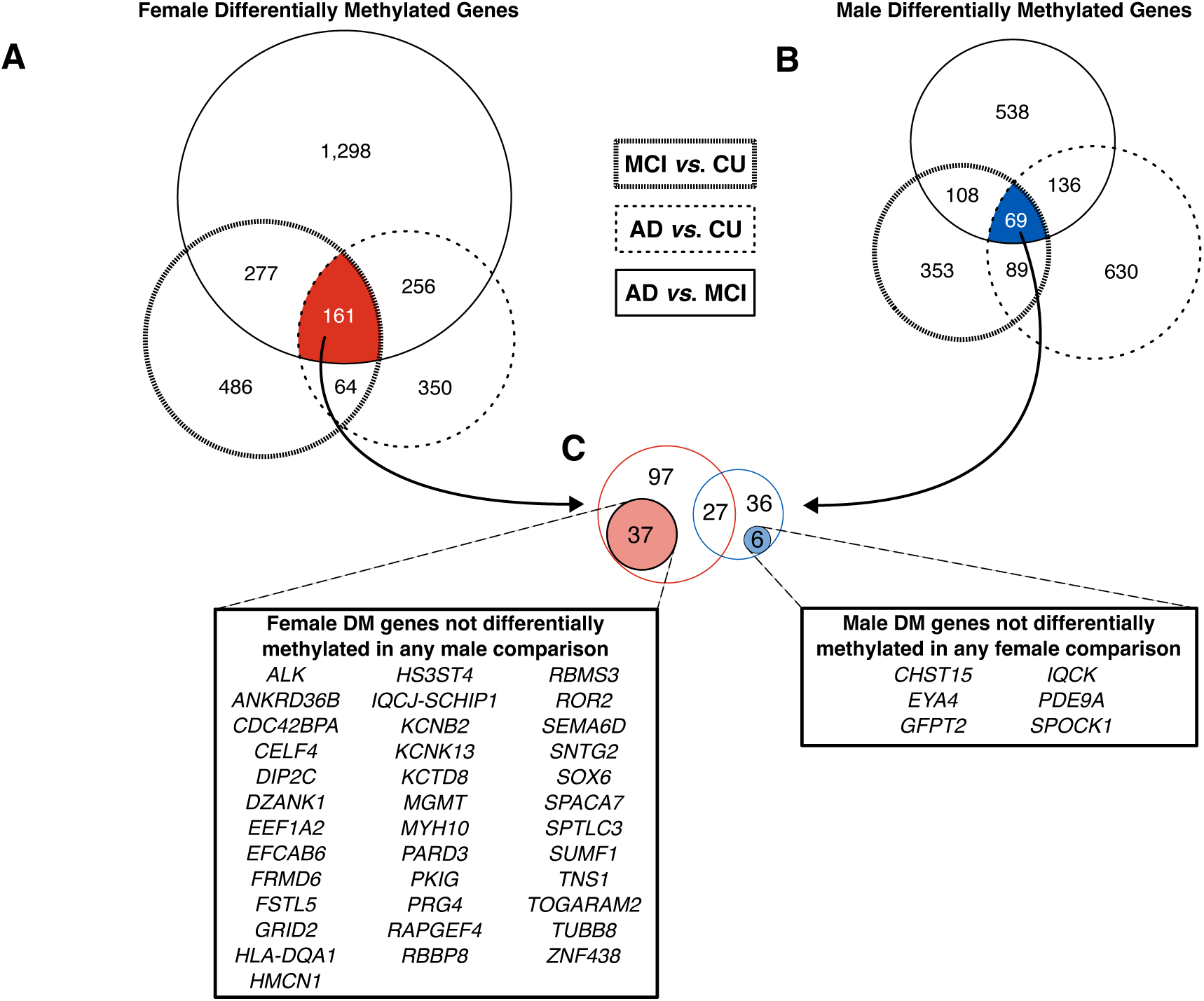
Sex-specific differentially methylated genes across comparisons. Venn diagram depicting the overlap of differentially methylated genes in the 3 female **(A)** and male **(B)** pairwise comparisons. (C) shows the Venn diagram of the overlap of the female (*N*=161) and male (*N*=69) differentially methylated genes. Inset circles with callouts identify genes that are differentially methylated in the 3 sex-specific comparisons but not differentially methylated in any of the opposite sex comparisons.

Gene ontological enrichment analysis of the 161 differentially methylated genes shared between all female pairwise comparisons yielded terms and biological processes related to synaptic organization and function including cell-cell adhesion via plasma membrane adhesion molecules, cell junction assembly, and regulation of synapse structure or activity (Supplemental Table 3), together with biological processes linked to neuron development including neuron differentiation, neuron projection development, and central nervous system development. Gene ontological enrichment analyses of the 69 differentially methylated genes shared between all male pairwise comparisons yielded no significantly enriched terms (Supplemental Table 3).

### Blood-specific enhancers and promoters comprise sex-specific DMPs

To test whether DMPs located >5 kilobases beyond a gene boundary (*i.e*., > 40% of all DMPs, Supplemental Fig. 1) comprise putative enhancer regions that interact with a gene promoter, we tested 58,654 recognized blood-specific enhancer-promoter sequences for the presence of female- and male DMPs in each pairwise comparison (Supplemental Table 4).

#### Sex-specific DMPs in blood-specific enhancers and promoters between MCI and CU

Analyses of enhancer regions identified 157 enhancers comprising 262 MCI-associated DMPs in females, and 97 enhancers comprising 171 MCI-associated DMPs in males. Filtering these data to DMPs residing in *cis* regulatory element (cCRE) enhancers^16^ identified 112/262 MCI-associated DMPs in females and 72/171 MCI-associated DMPs in males, including male-specific differential methylation of an enhancer of *CORO2A* that encodes a protein that participates in long term memory (Fig. 4A, Table 2). Analyses of promoter regions identified 82 promoters comprising 171 MCI-associated DMPs in females, and 70 promoters comprising 137 MCI-associated DMPs in males. Filtering these data to DMPs residing in cCRE promoters^16^ identified 8/171 MCI-associated DMPs in females and 6/137 MCI-associated DMPs in males.

**Figure 4:**
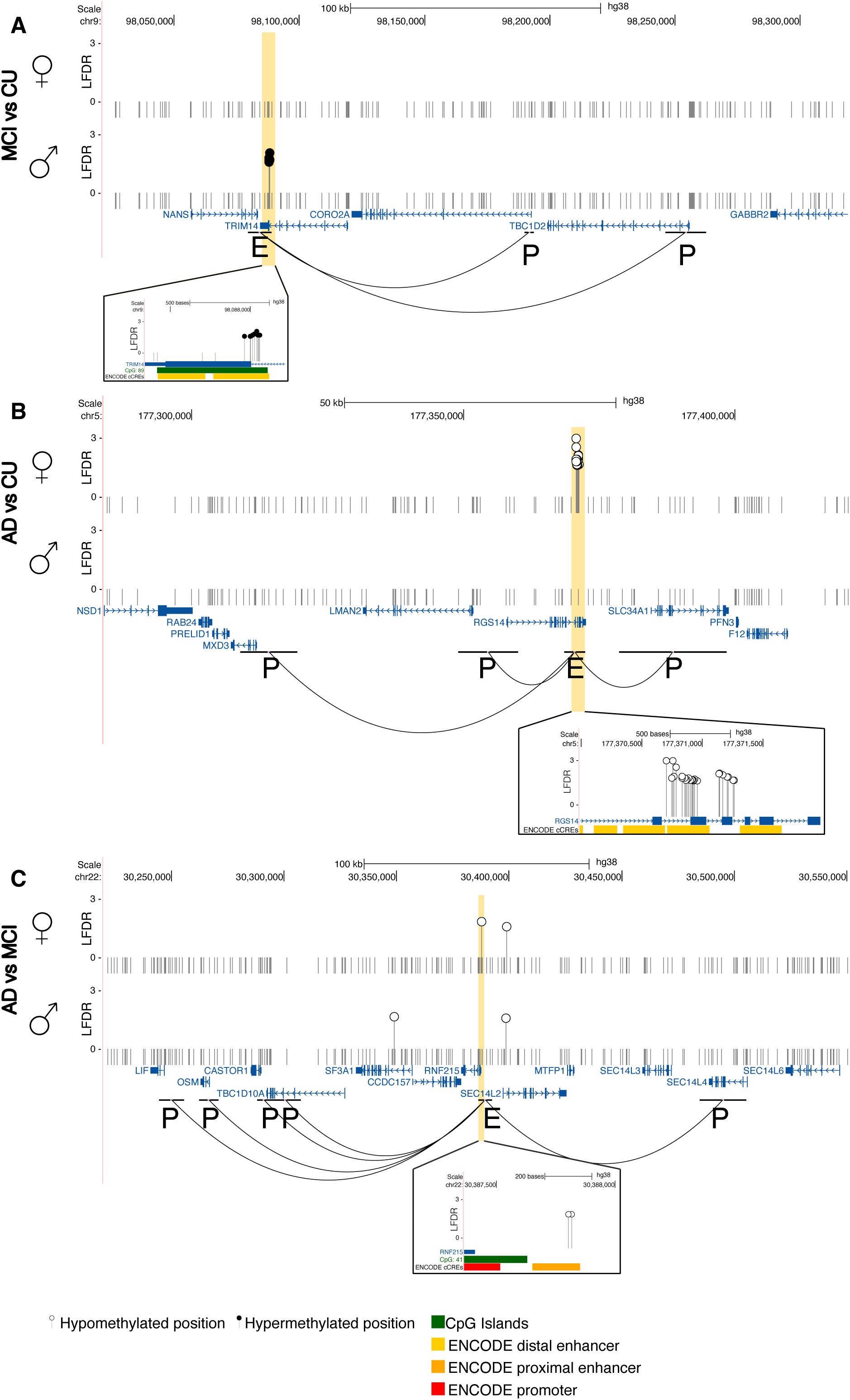
Long range promoter-enhancer interactions comprising DMPs. Long-range interactions with *CORO2A* and *TBC1D2* (antisense strand) in the MCI *vs.* CU comparison **(A)**. Long-range interactions with *MXD3, RGS14, LMAN2,* and *SLC34A1* in the AD *vs.* CU comparison **(B)**. Long-range enhancer interactions with *SEC13L4, CASTOR1, OSM,* and *LIF* in the AD *vs.* MCI comparison **(C)**. The relative scale (in kilobases [kb]) and genomic coordinates (in base pairs [bp]) of the region are provided at the top of panel. Significant differences in local false discovery rate (LFDR) adjusted DNA methylation *P*-values are provided (y-axis) with stems to indicate hypermethylation (black circles) or hypomethylation (white circles). One in 25 non-significant DMPs are depicted in each region (small stems with small gray circles). The relative location of all genes in the region is shown (bottom, blue) indicating the transcription direction (arrowheads). Enhancers (E) interact with the promoters (P) as depicted by arched loops. Zoomed-in regions (orange box) display annotated CpG islands (green) and ENCODE *cis* Regulatory Elements (cCREs) (red = promoter signature, orange = proximal enhancer signature, and yellow = distal enhancer signature).

#### Sex-specific DMPs in blood-specific enhancers and promoters between AD and CU

Analyses of enhancer regions identified 130 enhancers comprising 211 AD-associated DMPs in females, and 154 enhancers comprising 231 AD-associated DMPs in males. Filtering these data to DMPs residing in cCRE enhancers^16^ identified 90/211 AD-associated DMPs in females and 114/231 AD-associated DMPs in males, including female-specific differential methylation of an enhancer of *RGS14* that encodes a protein that participates long-term potentiation (Fig. 4B, Table 2). Analyses of promoter regions identified 52 promoters comprising 74 AD-associated DMPs in females, and 85 promoters comprising 145 AD-associated DMPs in males. Filtering these data to DMPs residing in cCRE promoters^16^ identified 14/74 AD-associated DMPs in females and 5/145 AD-associated DMPs in males.

#### Sex-specific DMPs in blood-specific enhancers and promoters between AD and MCI

Analyses of enhancer regions identified 362 enhancers comprising 448 DMPs between MCI and AD in females, and 105 enhancers comprising 138 DMPs between MCI and AD in males. Filtering these data to DMPs residing in cCRE enhancers^16^ between AD and MCI identified 153/448 DMPs in females and 37/138 DMPs in males, including female-specific differential methylation of an enhancer of *OSM* that encodes a protein regulator of α_1_-Antichymotrypsin linked to the progression of dementia (Fig. 4C, Table 2). Analyses of promoter regions between AD and MCI identified 158 promoters comprising 237 DMPs in females and 65 promoters comprising 115 DMPs in males. Filtering these data to DMPs residing in cCRE promoters^16^ identified 14/237 DMPs in females and 0/115 DMPs in males.

### Sex-specific differential methylation of transcription factor binding motifs between pairwise comparisons of cognitive status

Twenty five percent of MCI and AD sex-specific DMPs reside in promoter regions (*i.e*., 1-5kb upstream of the TSS and its putative promoter, Supplemental Fig. 1), suggesting that they may play a role in transcription factor binding. Accordingly, we tested for sequence motif enrichment within DNA sequences flanking DMPs in females and males (Figs. 5A and 5B; Supplemental Table 5). DNA sequences flanking DMPs between MCI *vs*. CU were enriched for transcription factor binding motifs in females (*N*=94) and males (*N*=59; E-value < 0.01). DNA sequences flanking DMPs between AD *vs*. CU were enriched for transcription factor binding motifs in females (*N*=66) and males (*N=*82; E-value < 0.01). DNA sequences flanking DMPs between AD and MCI were enriched for transcription factor binding motifs in females (*N*=120) and males (*N*=73; E-value < 0.01). Shared transcription factor binding motifs were found between the 3 female and 3 male pairwise comparisons (females *N*=61, males *N*=43, Fig. 5C). An enrichment of 37 differentially methylated transcription factor binding motifs were observed in both sexes between the 6 pairwise comparisons, indicating that regulatory elements in disease-associated methylation changes are shared between the sexes.

**Figure 5:**
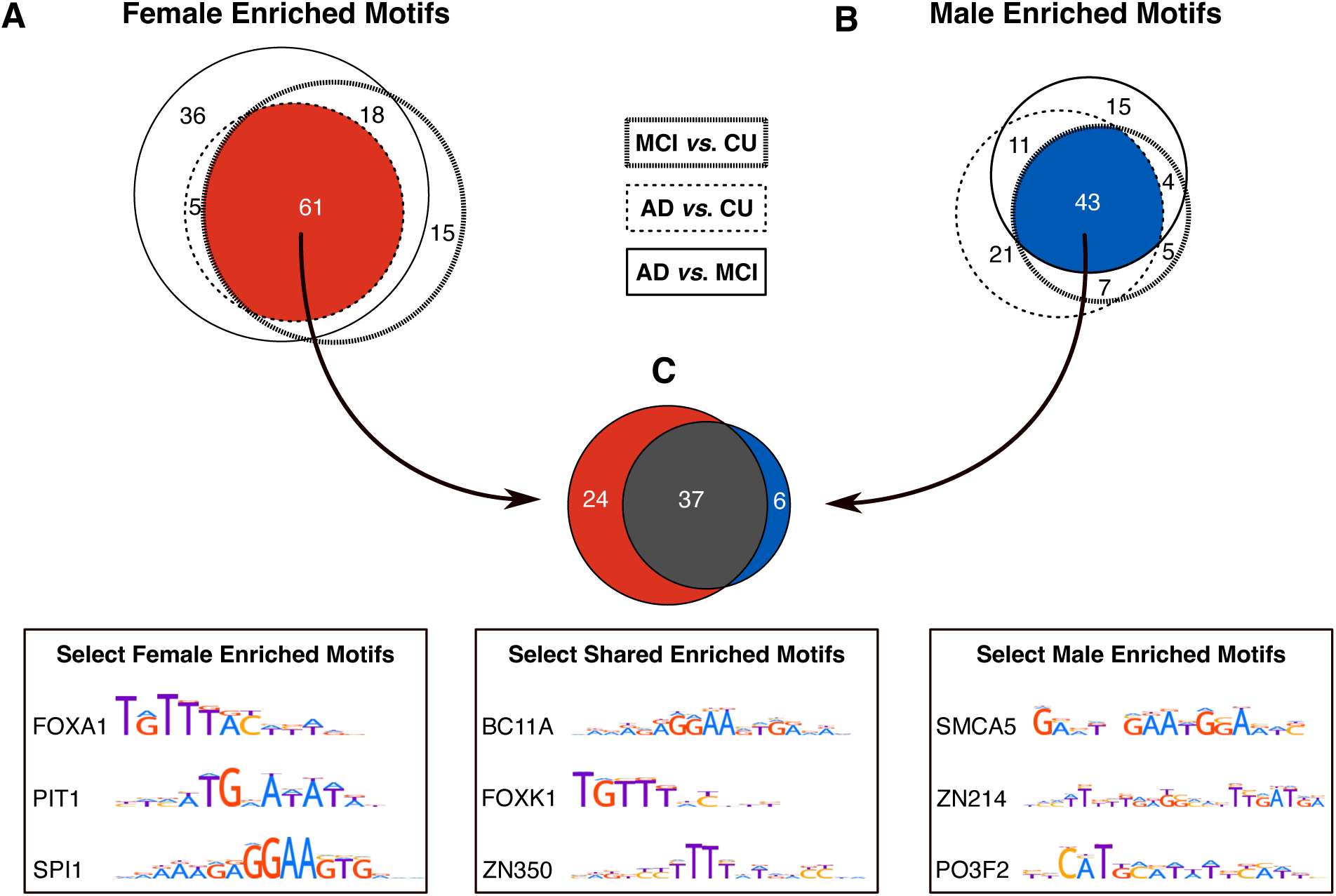
Transcription factor motif enrichment comprising DMPs. Venn diagram of the overlap of enriched transcription factor motifs for DMPs for females **(A)** and males **(B). (C)** Shows a Venn diagram of the overlap of female-specific and male-specific transcription factor motifs. Logo plots of representative transcription factor binding motifs comprising DMPs enriched in females (*N*=24), shared between the sexes (*N*=37), and in males (*N*=6). Consensus logo plots and motif symbols are from the HOCOMOCO v11 core HUMAN dataset (see methods).

### Progressive increases and decreases in sex-specific DNA methylation levels of DMPs

To test whether DNA methylation levels change progressively at specific DMPs in tandem with cognitive status, we tested for DMPs with increasing or decreasing DNA methylation levels across the three cohorts and found 16 female-specific DMPs and 15 male-specific DMPs (Table 3, Supplementary Table 6). 12 female-specific DMPs and 11 male-specific DMPs showed decrements in methylation levels across cognitive statuses (LFDR < 0.05, Fig. 6A and 6C), while 4 female-specific DMPs and 4 male-specific DMPs showed increments in methylation levels (LFDR < 0.05, Fig. 6B and 6D). Several of these sex-specific DMPs that change progressively in tandem with cognitive status overlap with transcription factor binding sites in blood (Table 3).^17^ These data indicate that blood DNA methylation levels may anticipate changes in cognitive status due to AD.

**Figure 6:**
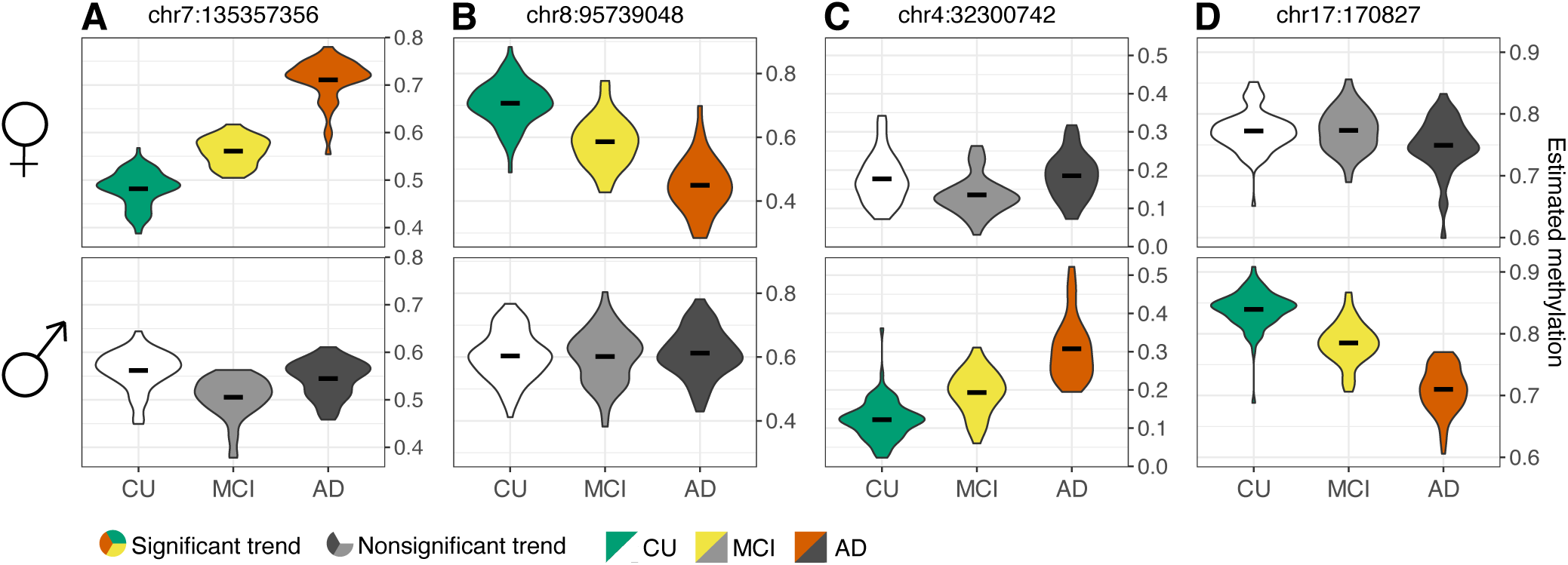
Progressively increasing or decreasing methylation across cognitive status. Violin plots depicting the estimated methylation values in females (top) and males (bottom) of each CpG locus (hg38 coordinates listed at top) are shown with the diagnosis (*x*-axis) and the percent methylation level (*y*-axis) for differentially methylated positions (DMPs) with significant (colored) and non-significant (grayscale) increments and decrements in DNA methylation levels among cognitively unimpaired (CU, green) persons and those with mild cognitive impairment (MCI, yellow) and Alzheimer’s disease (AD, orange).

**Table 3:**
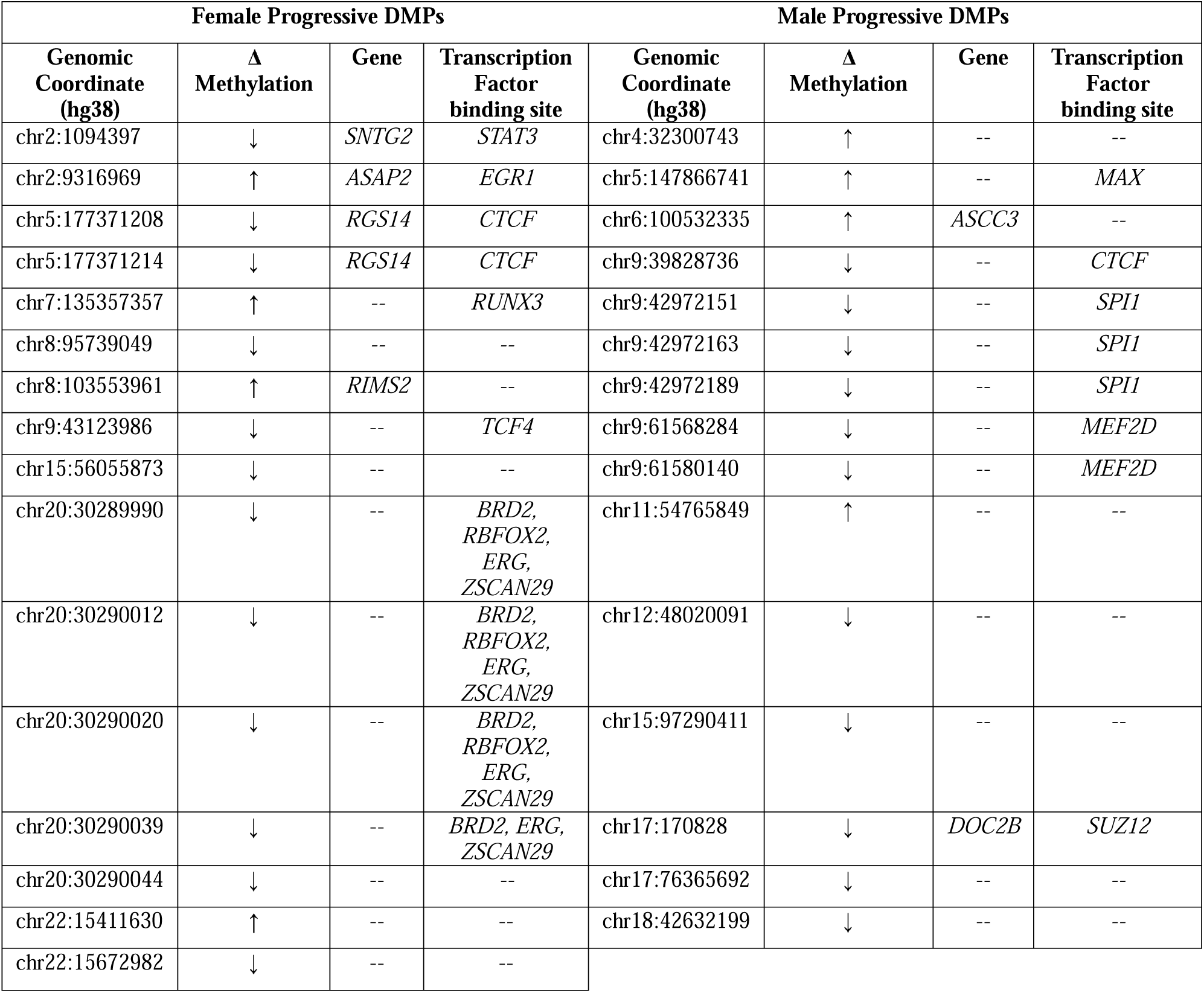
Progressive DMPs across cognitive status.

## Discussion

Blood DNA comprises over 6,200 sex-specific differentially methylated positions (DMPs) and over 1,200 sex-specific differentially methylated genes between persons who are cognitively unimpaired (CU) and those with mild cognitive impairment (MCI) and late onset dementia due to Alzheimer’s disease (AD) with minimal overlap between the sexes. These data indicate that levels and patterns of DNA methylation in blood are markedly distinct between females and males with MCI and AD. Finding that differentially methylated genes in blood are observed in dementia-related biological processes in the brain including regulation of membrane potential and synapse organization supports blood DNA methylation levels as candidate sex-specific biomarkers of AD susceptibility, diagnosis, and prognosis.

Multiple sex-specific differentially methylated genes observed in all 3 female pairwise comparisons (and not in any male comparison) participate in pathways with recognized links to AD including *GRID2* that encodes a candidate ionotropic receptor that participates in glutamate transfer and facilitates cerebellar synaptic organization and transmission (Fig. 3, Table 2). The protein encoded by *SPTLC3* supports cognitive performance with brain atrophy in AD (Table 2). The protein encoded by *ALK* mediates tau proteinopathy and disrupts autophagosome maturation and with tau accumulation and aggregation leading to neuronal dysfunction in AD (Table 2). Multiple sex-specific differentially methylated genes observed in all 3 male pairwise comparisons (and not any female comparison) participate in pathways with recognized links to AD including the *GFPT2* encoded protein that regulates glutamate metabolism and glucose control that underlies hypometabolism observed in AD (Fig. 3, Table 2). Variations in *EYA4* expression increase susceptibility to tau pathology in AD (Table 2). The protein encoded by *SPOCK1* is a member of the Sparc family that maintains the integrity of the blood-brain barrier. SPOCK1 also modulates the endocytic pathway of APP, a precursor to amyloid-β (Table 2).

Sex-specific differentially methylated enhancer regions comprising clusters of DMPs may disrupt recognized interactions with a gene promoter. For example, 17 of 25 AD-associated female-specific DMPs are observed in ENCODE *cis* regulatory element (cCRE) enhancer regions within the gene encoding *RGS14*, a GTPase-activating member of the regulator of G-protein signaling family of proteins highly expressed in the hippocampus participating in the processing of memory-related information (Table 2). This enhancer region interacts with promoter regions of multiple genes including *RGS14*, *LMAN2*, *MXD3*, and *SLC34A1*. Altered *RGS14* expression in rats and mice modifies long-term potentiation of memory.^18^

Because complex human disorders disrupt transcription factor (TF) function we tested for enrichments of sex-specific TF binding motifs that comprise DMPs in persons in our data.^19^ An enrichment of the *FOXA1* binding motif was differentially methylated in females with MCI and AD. Alterations of the *FOXA1* regulatory network are closely associated with AD (Table 2). An enrichment of the PO3F2 binding motif that participates in neuronal differentiation was differentially methylated in males with MCI and AD (Table 2). Thirty-seven differentially methylated TF binding motifs are shared between both sexes with MCI and AD, including BC11A that participates in differentiation of cortical neurons and is downregulated in AD (Table 2).

Few previous reports describe differential sex-specific DNA methylation levels in persons with MCI and AD.^20–22^ All are constrained by use of array-based platforms that interrogate less than 4% of the over 25,000,000 CpGs that we tested using whole genome methylation sequencing (WGMS). Controlling for sex as a covariate precludes identification of sex-specific findings. The present WGMS data overlap with DMPs in 3 genes from array-based reports: 1) Maturin, the neural progenitor differentiation regulator homolog (*MTURN*) is required for differentiation during primary neurogenesis and involved in the negative regulation of NF-kappaB transcription factor activity;^23^ 2) CDC42 binding protein kinase beta (*CDC42BPB*), a member of the serine/threonine kinase family;^24^ and 3) ArfGAP with coiled-coil, ankyrin repeat and PH domains 1 (*ACAP1*) a GTPase activating transport protein.^25^ Over 80% of AD-related DMPs are greater than 4 kilobases away from a CpG island thereby highlighting the advantages of differential DNA methylation detection with WGMS rather than commercially targeted microarray-based platforms.

The present investigation has strengths and limitations. Because samples were acquired in a cross-sectional experimental design, a longitudinal experimental design with samples from the same participants progressing from CU to MCI to AD are a top priority for future studies. The sex-specific contributions from the X and Y chromosomes were excluded from analyses in keeping with prior EWAS studies because of copy number differences of the sex chromosomes between sexes and X chromosome inactivation in females that generates a trimodal percent methylation distribution. Present data standing alone cannot resolve sex-specific changes in DNA methylation levels that cause changes in cognitive status from those that are compensatory or consequential. WGMS data herein was generated from bulk populations of blood cells thereby compelling future studies of specific cell types using single-cell techniques including fluorescence-activated single nuclei sorting. Lastly, differential DNA methylation levels that may correspond to discordant pathogenic pathways in females and males require comparisons between blood and brain samples.

Strengths of the present study include clinical diagnoses of CU, MCI, and AD phenotypes in the WRAP and WADRC cohorts that are based on stringent, well-validated and consensual clinical and psychometric criteria. Sample sizes are substantial allowing us to perform sex-specific differential analyses with robust adjustment for known confounders and multiple testing corrections. WGMS with comprehensive resolution of single CpG loci in the human genome using sequencing-by-synthesis is superior to partial coverage of hybridization-based microarray methods with loci and probes selected for commercial utility (∼4% of CpGs in the genome).

In conclusion, differentially methylated DNA positions in blood between persons who are cognitively unimpaired and those with MCI or AD are unexpectedly sex-specific and are identified within protein coding genes in biological pathways that are distinct between females and males. Sex-specific DNA methylation blood-based biomarkers that distinguish cognitive status will enhance the utility and precision of conventional neuro-imaging and plasma AD biomarkers.

## Methods

### Study Participants

This research was conducted in accord with the Declaration of Helsinki. The experimental protocol was approved by the institutional review board (IRB) of the University of Wisconsin School of Medicine and Public Health, Madison, WI. Written consent for study participation was obtained from each participant. Participants were enrolled in the Wisconsin Alzheimer’s Disease Research Center (WADRC^26^), and the Wisconsin Registry for Alzheimer’s Prevention (WRAP^27^). Details of the study design and demographic characteristics have been previously published.^14,15^ Participants were classified as cognitively unimpaired (CU) or as meeting criteria for MCI or AD based on the National Institute on Aging-Alzheimer’s Association thresholds by consensus conference.^28,29^ Samples from participants whose clinical status reverted to CU, or who were diagnosed as having non-Alzheimer’s disease dementia at a subsequent visit, were excluded.

### DNA extraction and generation of whole genome methylation sequencing (WGMS) data

Blood samples were acquired on the visit nearest following the date of MCI and AD diagnosis and matched for age by sample visit number in the CU cohort as previously described.^14^ High molecular weight genomic DNA was extracted, and sequence libraries were constructed using the NEBNext Enzymatic Methyl-seq (EM-seq™) kit (Ipswitch, MA) for whole genome methylation sequencing on an Illumina NovaSeq6000 sequencer (San Diego, CA). Image processing and sequence extraction used the Illumina Pipeline (Illumina, San Diego, CA).

### Data processing

Data processing was performed with the ENCODE gemBS-based processing pipeline as previously reported.^14^ Raw WGMS data were trimmed of adaptor sequences (Trim Galore) and aligned to the GRCh38.14 (hg38) human reference sequence. 5’ cytosines followed by a guanine (cytosine-phosphate-guanine; CpGs) are candidates for differential methylation to be identified as DMPs. CpG loci were filtered to have median coverage of 5 or greater with 50% or fewer missing values among all participants.

### Statistical analysis

#### Cell-type deconvolution

Blood samples comprise a composite of different white blood cell types each with a characteristic pattern of DNA methylation. To adjust for leukocyte heterogeneity, we used reference-free deconvolution of subpopulations existing in methylation data (DXM) to estimate the constituent fraction of each of the white blood cell types.^30^

#### Differential methylation analysis

To detect potential batch effects between the WADRC and WRAP cohorts, age, and sex (Table 1) principal components (PC) were calculated on 5% of CpGs with the greatest DNA methylation variability on each chromosome. Sequence data was imported into R (v4.4.0). Dispersion shrinkage for sequencing (DSS)^31^ was used to fit a beta-binomial model for each CpG locus adjusted for estimated white blood cell proportions, the first 2 PCs, age, and body mass index (BMI). All 382 samples were used for model fitting. A cognitive status sex-specific effect on methylation level for each CpG was interrogated with a Wald test of the corresponding regression coefficient comparing 6 pairwise groups of interest (*i.e*., female MCI *vs*. female CU, female AD *vs*. female MCI, female AD *vs*. female CU and 3 parallel male pairwise comparisons). The empirical distribution of the 25,406,945 *P*-values resulting from the theoretical null (standard normal distribution) for Wald tests from each comparison deviated from the expected uniform distribution with enrichment around small *P*-values. Accordingly, under-dispersed theoretical null test statistics were used to model the empirical null and to calculate local false discovery rates (LFDRs) with fdrtool^32^ and correct for multiple testing. For pairwise comparisons of sex-cognitive status (*e.g*., female AD *vs.* female CU, *etc.*), a CpG locus with a LFDR < 0.05 and an estimated mean methylation difference in sex-cognitive status group >5% was identified as a DMP.

#### Gene annotation, gene ontology analysis, and gene structure identification

To delimit gene sequence boundaries, 5′ to 3′ genomic coordinates were obtained from ENSEMBL (v111). Genes were defined as DNA sequences spanning 3 kilobases (kb) 5′ of a transcription start site (TSS) to 200 base pairs (bp) 3′ of a transcription termination site (TTS). Two criteria were used to identify differential methylation in genes. First, each gene must comprise a gene-wide LFDR < 0.05. To calculate gene-wide LFDRs, *P-*values of each CpG within the gene are aggregated into a harmonic mean *P*-value.^33^ The harmonic mean *P*-value controls the family-wise error rate of multiple statistical tests and does not assume independence among tests. The harmonic mean *P*-value thereby provides a preferred framework for aggregating *P*-values to correct for gene size (*i.e*., number of CpGs), and for highly correlated methylation levels of neighboring CpGs. This step was repeated for each gene yielding a single *P*-value for the whole gene. LFDRs were then calculated from the *P*-values to provide gene-wide false discovery rates. The second criterion to test for differential methylation of genes required at least one DMP defined as a CpG with an LFDR < 0.05 within the gene and a mean methylation difference of > 5%. Of note, this annotation may assign a DMP to more than a single gene. Gene Ontology (GO) analyses were performed with clusterProfiler (version 4.12.6),^34^ that conducts an over-representation test.^34,35^ False discovery rates were computed from *P*-values to correct for multiple testing. Sex-specific DMPs and differentially methylated genes in the present investigation do not identically align with those previously reported by Madrid *et al.* 2025^14^ in which sex was treated as a covariate rather than as an independent variable, with differences in sex-specific imputations for missing data and more stringent sex-specific effect size thresholds in the present data.

#### Differential methylation analysis of blood promoter–enhancer interactions

To identify differential methylation in gene promoter and enhancer regions in groups that differ by sex and cognitive status, promoter–enhancer interactions in blood cells were examined using published Promoter Capture Hi-C (PCHi-C) sequencing data.^36^ Interactions were filtered to have a median CHiCAGO score > 5 across all blood cell types to ensure adequate signal. After removal of false positive (*e.g*., bait–bait) and sex chromosome interactions, remaining promoter/enhancer interactions were lifted over to the human genome version GRCh38.14 with the UCSC liftOver tool. Interactions were filtered to remove any promoter or enhancer bin that spanned larger than 30kb. This approach retained 104,015 promoter–enhancer interactions comprising 9,507 distinct promoters and 58,430 distinct enhancers for comparison to the WGMS data. Promoters and enhancers were considered differentially methylated if they contained one or more DMPs. DMPs located within interacting enhancer/promoter regions were further filtered for ENCODE candidate *cis* regulatory elements (cCREs).^16^

#### Motif Enrichment analysis

To investigate regulatory mechanisms associated with sex-specific differential DNA methylation in MCI and AD, we performed motif enrichment analysis (MEA) on DMPs that differ by sex and cognitive status using the Analysis of Motif Enrichment (AME) tool from the MEME suite.^37^ Six browser extensible data (BED) files were generated by adding 25 flanking base pairs to each DMP in the 6 pairwise comparison DMP lists. A background BED file was generated by randomly selecting 500,000 CpG loci (out of 25,406,945 CpGs). BED file regions were then merged within each list if they overlapped or were directly adjacent to each other (bedtools, v2.31.1). The resulting BED file regions were transferred to AME software (v5.5.7, https://meme-suite.org/meme/tools/ame). Non-default options included: (1) background control sequences comprising the background BED file (used for 6 enrichment analyses) and (2) motif database HOCOMOCOv11 core HUMAN.^38^ Enriched motifs of recognized human transcription factors (*N*=401) were identified using a Fischer’s exact test (AME default) and an *E*-value < 0.01 was adopted for all comparisons.

#### DNA methylation level trends across sex and cohort cognitive status

DSS was used to identify CpG positions with linear methylation trends across the 3 cognitive statuses (*i.e.*, CU, MCI, AD) in females and males. A CpG locus with an LFDR threshold of < 0.05, coupled with > 5% methylation difference between cognitive status groups was identified as a progressive DMP. Progressive DMPs were queried against the ChIP-Atlas^17^ to assess their overlap with transcription factor binding sites identified by ChiP-seq in blood utilizing the peak browser online tool.^39^ DMPs were queried against the “ChIP: TF and others” (*track type class*) and “blood” (*cell type class*). The top scoring transcription factor peak was reported in Table 3 (multiple transcription factors were reported if they all had maximum peak scores of 1000).

## Data Availability

All processed sequencing data generated in this study will be submitted to dbGAP upon publication.

## Acknowledgements

The authors would like to acknowledge and thank the participants and study personnel who made this work possible. This work was supported by the National Institutes of Health grant numbers: R01AG066179; R01AG021155; R01AG027161; P30AG062715; and HG003747, as well as the Alzheimer’s Association AARF19-614533 and the UW Department of Neurological Surgery. PB was supported by the National Institute of General Medical Sciences award number T32GM007133. The funding sources had no role in the conceptualization, design, data collection, analysis, decision to publish, or preparation of the manuscript.

## Authors’ Contributions

PEB, LAP, KJH, RSA, contributed to the conceptualization of the investigation. LRC, SA, SCJ provided resources. LAP provided methodology and data curation. PEB, AM, LAP, SK, KJH, RSA provided formal analysis and interpretation. PEB, LAP, KJH, RSA wrote the original draft. All authors provided review of early drafts of the manuscript.

## Ethics approval and consent to participate

The experimental protocol was approved by the Institutional Review Board (IRB) of the University of Wisconsin School of Medicine and Public Health, Madison, WI. All participants signed an IRB-approved informed consent.

