## Supplemental Figure 1 for "Sex-Specific Differential DNA Methylation in Mild Cognitive Impairment and Alzheimer’s Disease"

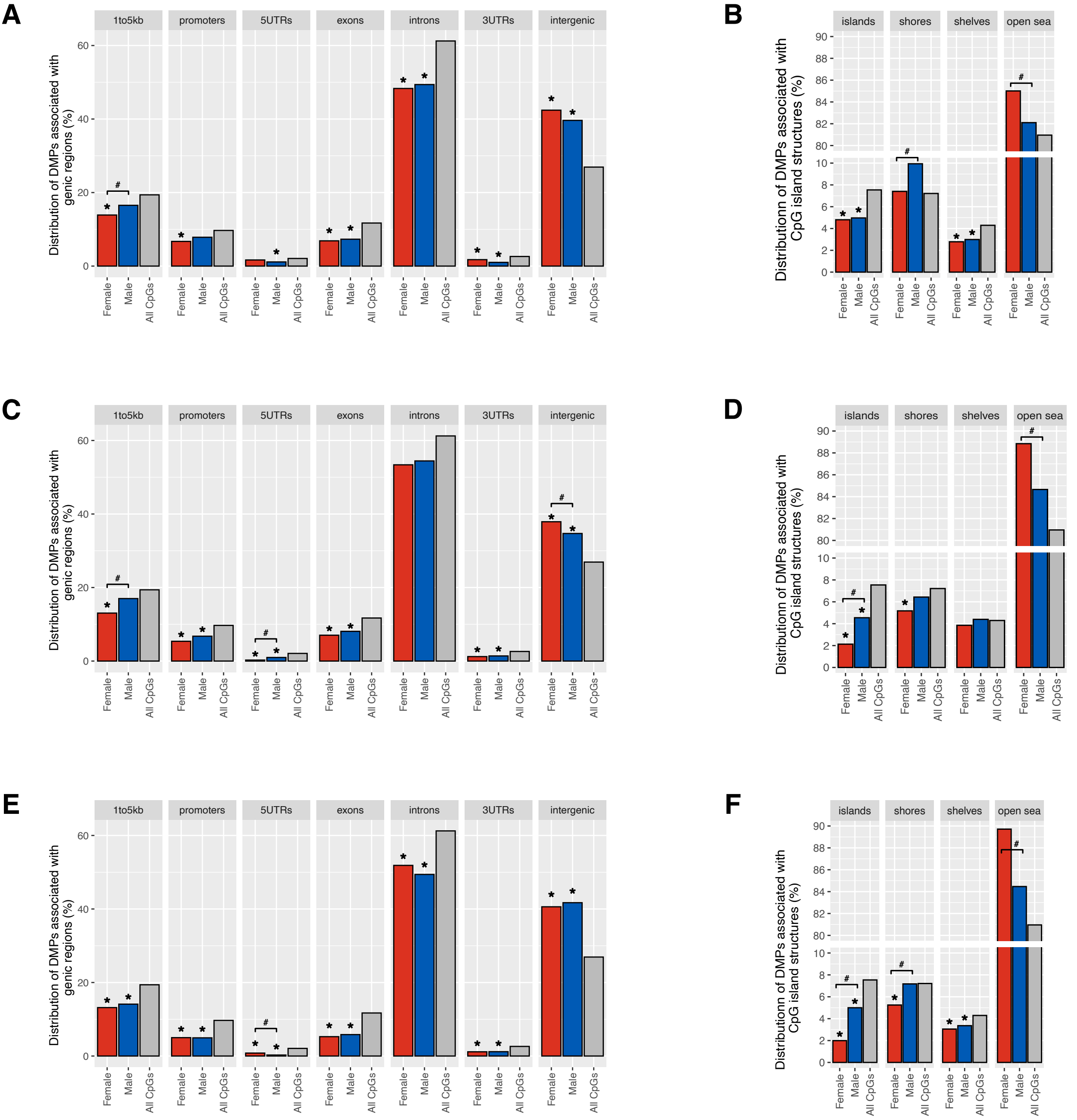


**Supplemental Figure 1:** Genic Structure bar plots depict annotation of both female- (red) and male-specific (blue) DMPs to genic structures (i.e., 1-5kb upstream, promoters [1kb upstream to transcription start site], 5' untranslated region, exons, introns, 3' untranslated region, and intergenic) for the 3 comparisons (**A**) MCI vs CU, (**C**) AD vs CU, and (**E**) AD vs MCI. Gray bars depict overall genic structure distribution of all CpG loci analyzed. * indicates significance of either the male or female genic region to the overall distribution (hypergeometric test, P-value < 0.01). # indicates significant differences between males and females (Fisher's exact test, P-value < 0.01). D) CpG Island bar plots depict the female (red) and male (blue) percentage of DMPs annotated to CpG islands, shores (2kb from island), shelves (2-4kb from island), and the open sea (>4kb from island) for the 3 comparisons (**B**) MCI vs CU, (**D**) AD vs CU, and (**F**) AD vs MCI. Gray bars depict overall (genomic) distribution of all CpG loci analyzed. * indicates significance of either the male or female genic region to the overall distribution (hypergeometric test, P-value < 0.01). # indicates significant differences between males and females (Fisher's exact test, P-value < 0.01).
