## Supplemental Figure 2 for "Sex-Specific Differential DNA Methylation in Mild Cognitive Impairment and Alzheimer’s Disease"

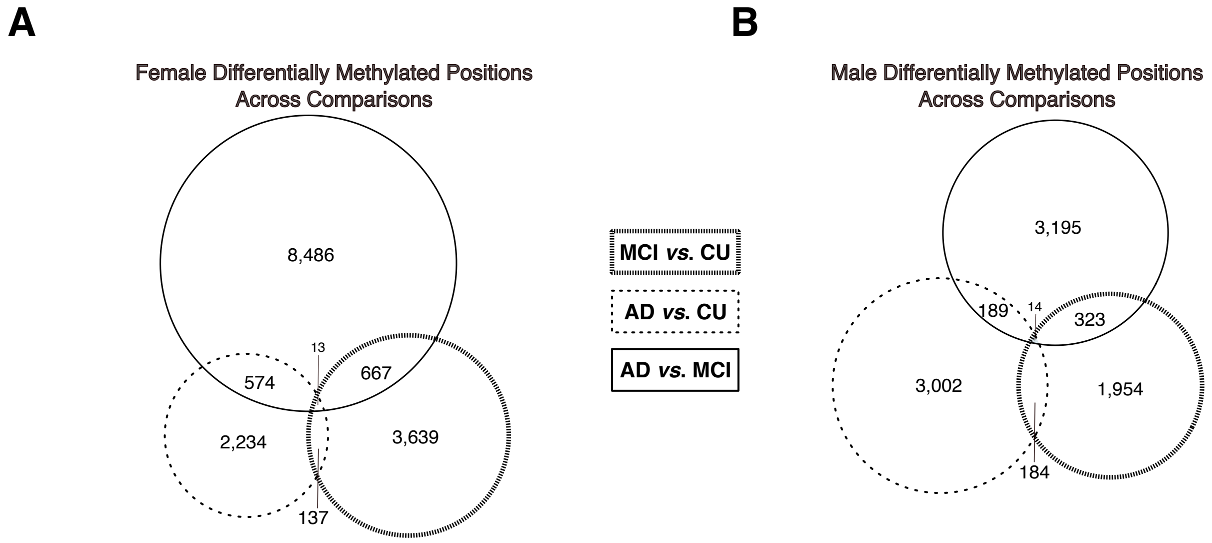


**Supplemental Figure 2:** Sex-specific differentially methylated position (DMP) differences shared across comparisons, highlighting distinct and opposing methylation patterns in example genes. **A**) Venn diagram depicting the overlap of DMPs within females in the 3 pairwise comparisons. **B**) Venn diagram depicting the overlap of DMPs within males in the 3 pairwise comparisons.
